# On the robustness of inference of association with the gut microbiota in stool, swab and mucosal tissue samples

**DOI:** 10.1101/2021.02.04.429844

**Authors:** Shan Sun, Xiangzhu Zhu, Xiang Huang, Harvey J. Murff, Reid M. Ness, Douglas L. Seidner, Alicia Sorgen, Ivory Blakley, Chang Yu, Qi Dai, M. Andrea Azcarate-Peril, Martha J. Shrubsole, Anthony A. Fodor

**Author notes:** **Corresponding Author:** Anthony A. Fodor, Department of Bioinformatics and Genomics University of North Carolina at Charlotte 9331 Robert D. Snyder Rd, Room 361 Charlotte, NC 28223. These authors contributed equally to this work (Martha J. Shrubsole, Anthony A. Fodor).

## Abstract

The gut microbiota plays an important role in human health and disease. Stool, swab and mucosal tissue samples have been used in individual studies to survey the microbial community but the consequences of using these different sample types are not completely understood. We previously reported differences in microbial community composition with 16S rRNA amplicon sequencing between stool, swab and mucosal tissue samples. Here, we extended the previous study to a larger cohort and performed shotgun metagenome sequencing of 1,397 stool, swab and mucosal tissue samples from 240 participants. Consistent with previous results, taxonomic composition of stool and swab samples was distinct, but still more similar to each other than mucosal tissue samples, which had a substantially different community composition, characterized by a high relative abundance of the mucus metabolizers *Bacteroides* and *Subdoligranulum,* as well as bacteria with higher tolerance for oxidative stress such as *Escherichia*. As has been previously reported, functional profiles were more uniform across sample types than taxonomic profiles with differences between stool and swab samples smaller, but mucosal tissue samples remained distinct from the other two types. When the taxonomic and functional profiles of different sample types were used for inference in association with host phenotypes of age, sex, body mass index (BMI), antibiotics or non-steroidal anti-inflammatory drugs (NSAIDs) use, hypothesis testing using either stool or swab gave broadly similar results, but inference performed on mucosal tissue samples gave results that were generally less consistent with either stool or swab. Our study represents an important resource for the experimental design of studies aimed to understand microbiota perturbations specific to defined micro niches within the human intestinal tract.

## Introduction

A growing number of studies have reported the essential roles of the human gut microbiota in human health and that microbiota alterations are associated with diseases including colorectal cancer, inflammatory bowel disease, obesity and diabetes [1–4]. The human colorectum is a complex system consisting of many microhabitats; studies have reported that the luminal and mucosal microbiota harbor heterogeneous microbial communities [5]. With the oxygen decline from the intestinal mucosa towards the lumen, anaerobic microorganisms are likely more abundant in luminal than mucosal environments [6]. On the other hand, the mucosal microbiota, directly adherent to the host tissue, may be more sensitive and respond more rapidly to localized changes in host tissues, compared to the luminal microbiota that is isolated from the loose mucus layer on the surface of the colorectal wall [7].

Stool samples are the most common biospecimen used to assess composition and functionality of the human gut microbiota in human research because of the large amount of biomass and the feasibility of collection; however, stool-derived profiles are more representative of luminal microorganisms than of mucosa-associated microbes. Mucosal tissue biopsy better characterizes mucosa-associated microbes but is less frequently used because of the invasive nature and accompanying risk of the procedure. Rectal swab may be used when stool samples are not practical to obtain, for example in the intensive care unit, and may collect a combination of both luminal and mucosal communities [8]. While stool and mucosal samples are generally distinct, there are mixed findings on the similarity between stool and swab samples [9–11]. Thus, different biospecimen types may be needed to sample microorganisms residing in different niches or to reflect different physiological conditions. For example, a study on colitis-induced inflammation in mouse reported that microbial dysbiosis in the mucus layer was detected preceding colitis while changes in stool microbiota were detected post-colitis [7].

Most of the studies assessing the variation of microbiota profile by biospecimen type have focused on taxonomic composition characterized by 16S rRNA amplicon sequencing. Previous literature of observed variation using shotgun metagenomics is usually limited by the sample size, including our own previous study [8]. Compared to the 16S rRNA amplicon sequencing, shotgun metagenome sequencing utilizes total DNA instead of PCR products thus reducing the bias introduced during processing. Moreover, metagenome sequencing not only determines the taxonomic composition of the gut bacterial communities but also generates information about functional information. With the increasing application of shotgun metagenome sequencing in microbiota studies, a better understanding of the metagenome variation across biospecimen types will help investigators develop and interpret their experimental design.

In this study, we collected matched stool, rectal swab and mucosal tissue samples from 240 study participants at two time points, which resulted in 1,397 shotgun metagenomes. This is one of the largest studies comparing metagenomes of human stool, rectal swab and colorectal mucosal tissue samples. We estimated the biospecimen type variation of both metagenome taxonomy and functional pathways. We also assessed whether the associations between taxa/pathways and age, sex, body mass index (BMI), non-steroidal anti-inflammatory drugs (NSAIDs) use and antibiotics use were consistent across the different sample types.

## Methods

### Study Population and Biospecimen Collection

The Personalized Prevention of Colorectal Cancer Trial (PPCCT) was a double-blind, placebo-controlled, randomized clinical trial designed to test the interaction between a *TRPM7* genotype and reduction of the calcium/magnesium intake ratio via magnesium supplementation on colorectal carcinogenesis biomarkers. Study design and biospecimen collection have been previously described [8]. In brief, participants were randomized to receive for 12 weeks either a personalized dose of placebo (microcrystalline cellulose) or magnesium (magnesium glycinate). Inclusion criteria included aged 40-85, personal history of colorectal polyps, known *TRPM7* rs8042919 genotype, and daily intakes of calcium between 700-2000 mg/day and the ratio of calcium to magnesium of 2.6 or greater. Exclusion criteria included pregnancy, breastfeeding, use of medications that may interact with magnesium, or personal history of cancer, colon resection or colectomy, inflammatory bowel disease, organ transplantation, gastric bypass, chronic diarrhea, chronic renal disease, hepatic cirrhosis, chronic ischemic heart disease, or Type I diabetes. All study procedures were performed in accordance with relevant guidelines and regulations as approved by the Vanderbilt Institutional Review Board. The study is registered at ClinicalTrials.gov (NCT01105169).

Biospecimens were collected at home or in an in-person study visit at the beginning of the trial (baseline) and at the conclusion of the study 12 weeks later (mean 12.3 ± 1.03 weeks) [8]. Stool samples were collected by study participants at home using a white plastic collection container covering the toilet bowl, aliquoted by the participant into sterile cryovials, and stored in the home freezer until transport with an ice pack to the study visit. Stool was collected up to 3 days prior to the study visit. Rectal swabs and mucosal tissues were collected by the study physician at the study visits. Rectal swabs were collected by inserting a culturette swab through the anal canal, swabbing the distal rectal mucosa, and placing the swab into a cryovial. Rectal mucosal samples were collected through an anoscope using standard mucosal biopsy forceps and these samples were placed into separate storage vials. All three biospecimen types were frozen at −80□°C until use.

### DNA Isolation and Sequencing

Samples were transferred to a 2 ml tube containing 200 mg of ≤106 μm glass beads (Sigma, St. Louis, MO) and 0.3 ml of Qiagen ATL buffer (Valencia, CA), supplemented with lysozyme (20 mg/ml) (Thermo Fisher Scientific, Grand Island, NY). The suspension was incubated at 37°C for 1 h with occasional agitation. Subsequently the suspension was supplemented with 600IU of proteinase K and incubated at 60°C for 1 h. Finally, 0.3 ml of Qiagen AL buffer were added and a final incubation at 70°C for 10 minutes was carried out. Bead beating was then performed for 3 minutes in a Qiagen TissueLyser II at 30Hz. After a brief centrifugation, supernatants were transferred to a new tube containing 0.3 ml of ethanol. DNA was purified using a standard on-column purification method with Qiagen buffers AW1 and AW2 as washing agents and eluted in 10mM Tris (pH 8.0).

Whole-genome shotgun metagenomics (WGS) DNA sequencing was performed as previously described [8]. Briefly, 1 ng of genomic DNA was processed using the Nextera XT DNA Sample Preparation Kit (Illumina). Next, fragmented and tagged DNA was amplified using a limited-cycle PCR program. In this step index 1(i7) and index 2(i5) were added between the downstream bPCR adaptor and the core sequencing library adaptor, as well primer sequences required for cluster formation. The DNA library was purified using Agencourt^®^ AMPure^®^ XP Reagent. Each sample was quantified and normalized prior to pooling. The DNA library pool was loaded on the Illumina platform reagent cartridge and on the Illumina HiSeq instrument.

### Bioinformatics and Statistical Analyses

Sequencing output from the Illumina HiSeq4000 platform was converted to fastq format and demultiplexed using Illumina Bcl2Fastq 2.18.0.12. Quality control of the demultiplexed sequencing reads was verified by FastQC. Human genome contamination was removed from the shotgun metagenome sequencing reads with KneadData. The number of reads before and after removing human genome contamination is shown in Fig. S1. The taxonomic composition of the filtered reads was characterized with MetaPhlAn2 [12] while the functional pathways were annotated with HUMAnN2 against the UniRef database [13]. Unmapped reads were excluded from the following analyses. PCoA ordination was generated with Bray-Curtis dissimilarity based on genus composition and functional pathway abundance respectively with function ‘capscale’ in the R package ‘vegan’. The PERMANOVA test was performed with the function ‘adonis’ in the same package. For each individual genus or pathway, we built linear mixed effects models with the function ‘lme’ in R package ‘nlme’ with the aim of examining differences between the modes based on sample type variation. The genera and pathways with presence <10% in all samples were excluded to avoid spurious results and P-values were adjusted with the Benjamini-Hochberg method for multiple testing.

Model 1 was used to test the associations between the metagenome and biospecimen types (stool, swab or mucosal tissue). Model 1 was performed for each pair of sample types to get the direction of changes and adjusted for host factors.

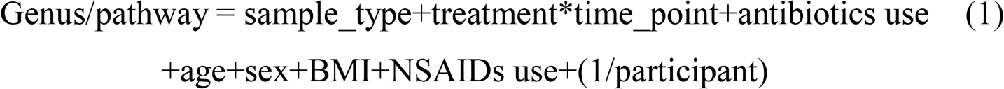

In this model, sample type, treatment, time point, age, sex, BMI, antibiotics and NSAIDs use were fixed effects while participant ID was a random effect. Using pairwise models allowed for direct comparison between sample types. The significance was determined as <10% FDRs corrected with Benjamini-Hochberg method. Significant genera and pathways identified in this model were plotted as heatmaps with the function ‘pheatmap’.

Model 2 was used to test the associations between metagenome and host factors in each sample type.

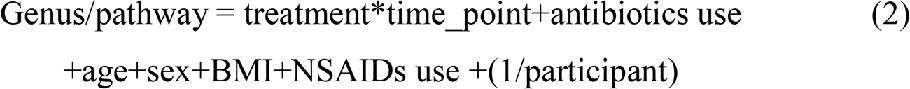

In this model, treatment, time point, age, sex, BMI, antibiotics and NSAIDs use were fixed effects while participant ID is a random effect. The correlations between inferences (−log10(P)) produced in different sample types were tested with Spearman correlations and the plots were generated with ‘ggplot2’.

Because of the compositional nature of the shotgun metagenome sequencing data, we also utilized ALDEx2 [14] which uses Bayesian methods and a geometric mean based normalization to minimize compositional artifacts to confirm our observations. Because ALDEx2 does not support models adjusted for covariates, the associations were tested with one variable models.

## Results

### Taxonomic composition of metagenomes was associated with sample types

After quality control, there were 1,397 stool, swab and mucosal tissue metagenomes from 240 participants. We characterized the taxonomic composition and functional pathways of the metagenomes and found substantial variation by sample type. Shannon diversity at the genus level was significantly different between sample types, with mucosal tissue samples of the lowest diversity and swab the highest (Fig. 1a). PCoA ordinations of genus composition showed a distinct cluster of mucosal tissue samples (Fig. 1b). A PCoA ordination in which mucosal tissue samples were excluded in order to better visualize the stool and swab samples showed clear separation as well (Fig. 1c). A PERMANOVA test indicated that the genus composition was significantly associated with sample type (P=0.001, with 999 permutations). The differences across stool, swab and mucosal tissue samples explained 31.6% of the variance, while the differences between stool and swab explained 5%, further supporting the observation that mucosal tissue samples were more distinct compared to stool and swab. Microbial taxonomic composition at other levels from phylum to species levels were also significantly associated with sample type (Table S1). The PERMANOVA tests and PCoA ordinations demonstrate that the microbial metagenomes sampled with different methods were different at the community level.

**Fig. 1.**
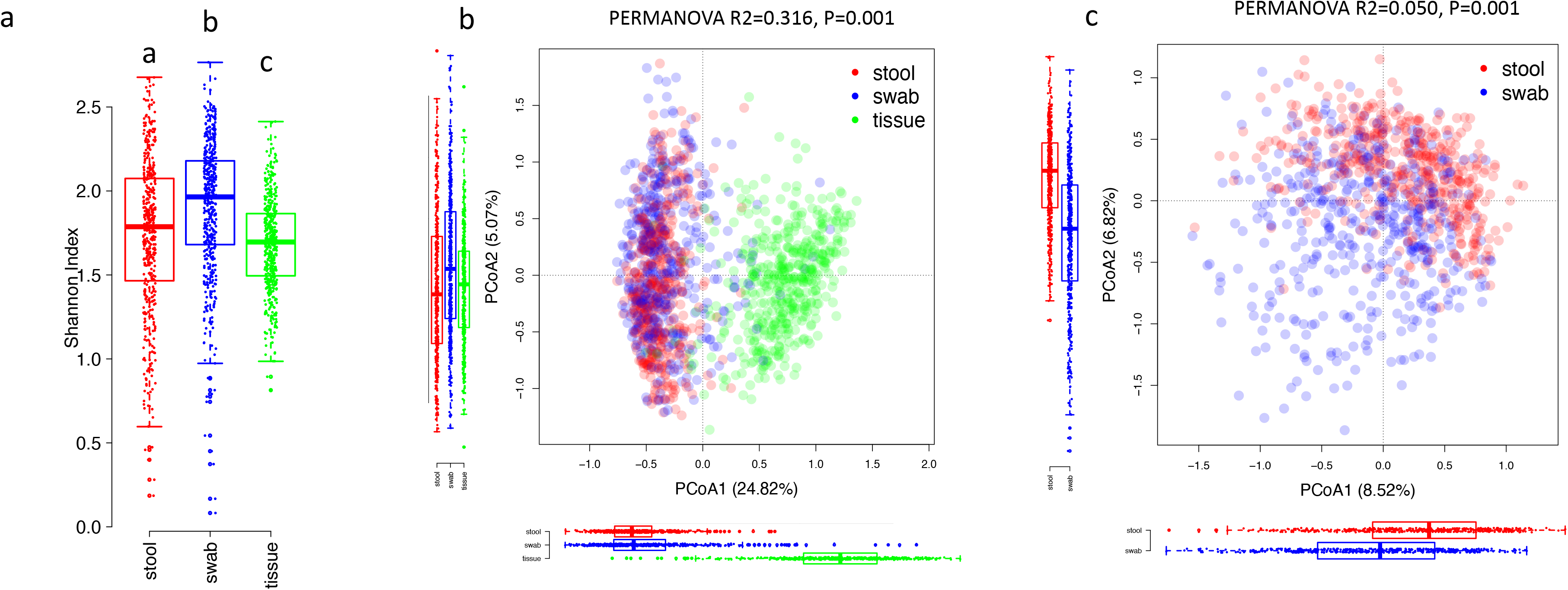
Alpha-diversity and PCoA ordinations of the taxonomic composition of microbial metagenomes at the genus level. Color indicates the sample types. (a) Alpha diversity across sample types. Differences between sample types were tested with Wilcoxon’s test. (b) Mucosal tissue samples formed a distinct cluster from stool and swab samples. (c) Separation of stool and swab samples.

In order to identify differentially abundant taxa, we used a linear mixed-effects models to compare the sample types in pairs (Model 1). Among the 60 genera with presence in >10% samples, 56 were different between at least one pair of samples, with 35 significantly different between stool and swab samples, 53 between stool and tissue, and 51 between swab and tissue (Fig. 2). Because the sequencing depths were different between sample types (Fig. S1), we also utilized an analysis pipeline based on ALDEx2, which attempts to explicitly correct for compositional artifacts. The differential abundance of the 56 taxa across sample types were supported by results from ALDEx2, except for *Paraprevotella* and an unknown genus of the Clostridiaceae family (Table S2). P-values from the two methods were generally consistent (Fig. S2a). Tissue samples had higher relative abundance of *Bacteroides*, *Subdoligranulum, Escherichia*, *Blautia* and unclassified genera of the families *Propionibacteriaceae* and *Acidaminococcaceae*. Compared to stool samples, swab samples were enriched in *Propionibacterium*, *Campylobacter*, *Porphyromonas*, *Prevotella*, *Clostridium*, *Streptococcus* and had lower abundance of *Methanobrevibacter*, *Dialister*, *Adlercreutzia*, *Haemophilus*, *Klebsiella*, *Akkermansia*, *Alistipes* and *Paraprevotella*.

**Fig. 2.**
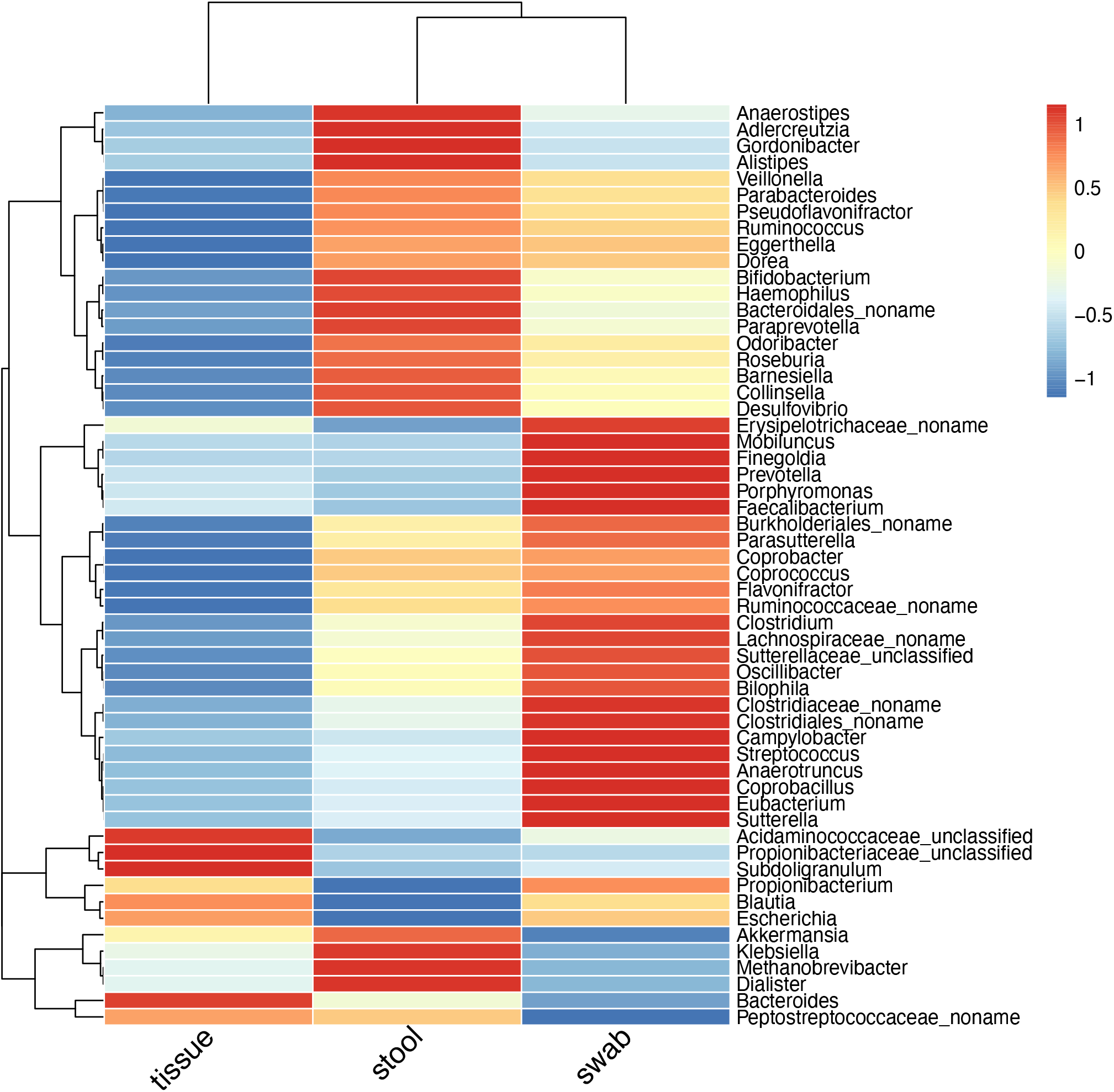
Heatmap of genera that were significantly different between sample types (FDR<0.05). Keys indicate the z-scores of averaged taxonomic abundance.

### Functional pathways of metagenomes were associated with sample types

The metagenomes of mucosal tissue samples had a higher number of reads that could not be mapped to the UniRef databases after removing host sequences (54% compared to 30% for stool and 28% for swab samples), indicating that the mucosal tissue microbiota was less represented in the current database. The number of microbial pathways was lower in mucosal tissues compared to other samples (Fig. 3a). The PCoA ordinations of functional pathways showed a similar specific cluster of mucosal tissue samples (Fig. 3b), while the stool and swab samples were less separated compared to the PCoA ordination based on genus composition (Fig.3c). A PERMANOVA test indicated that functional pathways were also significantly different across sample types (stool, swab and mucosal tissue: R^2^ = 0.273, P=0.001; stool and swab: R^2^ = 0.048, P=0.001). We again used a linear mixed effects model to identify the differential functional pathways between samples. In 343 functional pathways with presence in >10% samples, 318 were significantly different between at least one pair of samples, with 269 of differential abundance for stool-swab comparison, 222 for stool-tissue and 233 for swab-tissue (Fig. 4). Among the 318 significant pathways, only 8 were not supported by the analysis of ALDEx2 (Table S3; Fig. S2b).

**Fig. 3.**
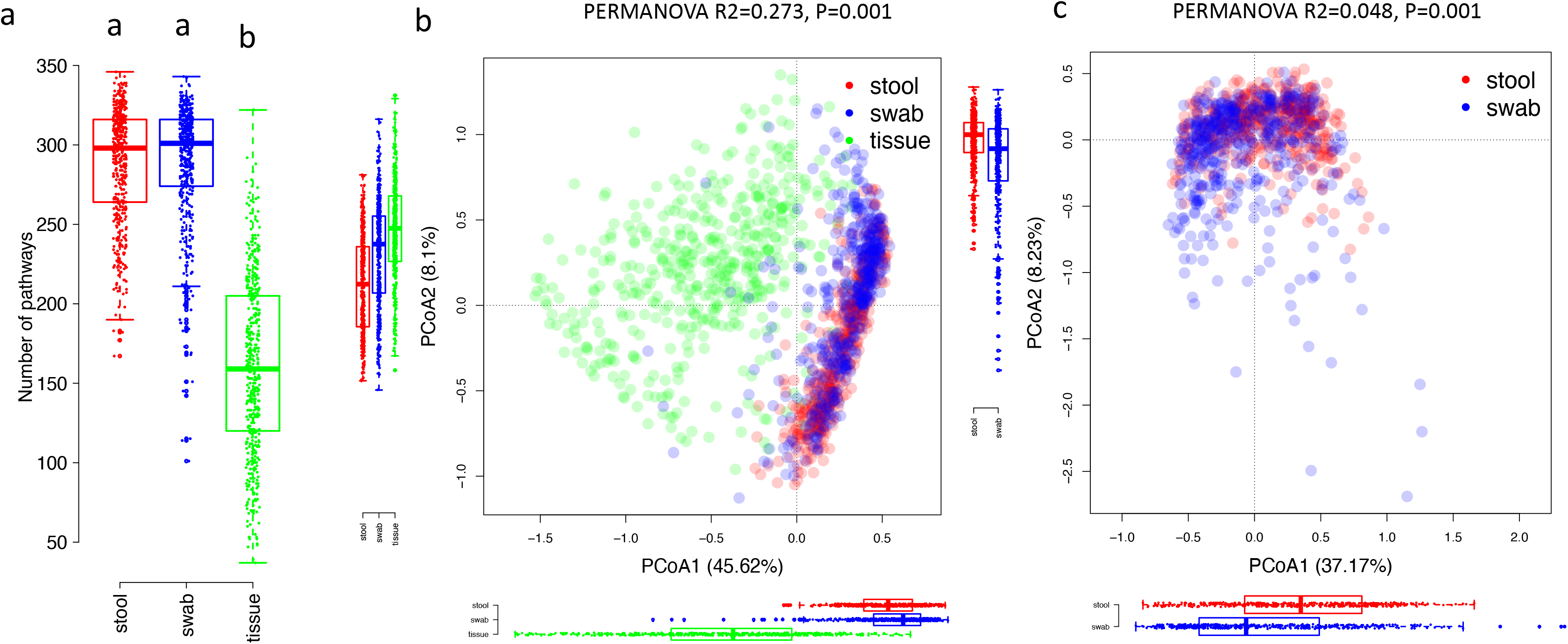
The number of pathways and PCoA ordinations of functional pathways of microbial metagenomes. Color indicates the sample types. (a) The number of pathways across samples. (b) Mucosal tissue samples formed a distinct cluster from stool and swab samples. (c) visualization of only stool and swab samples.

**Fig. 4.**
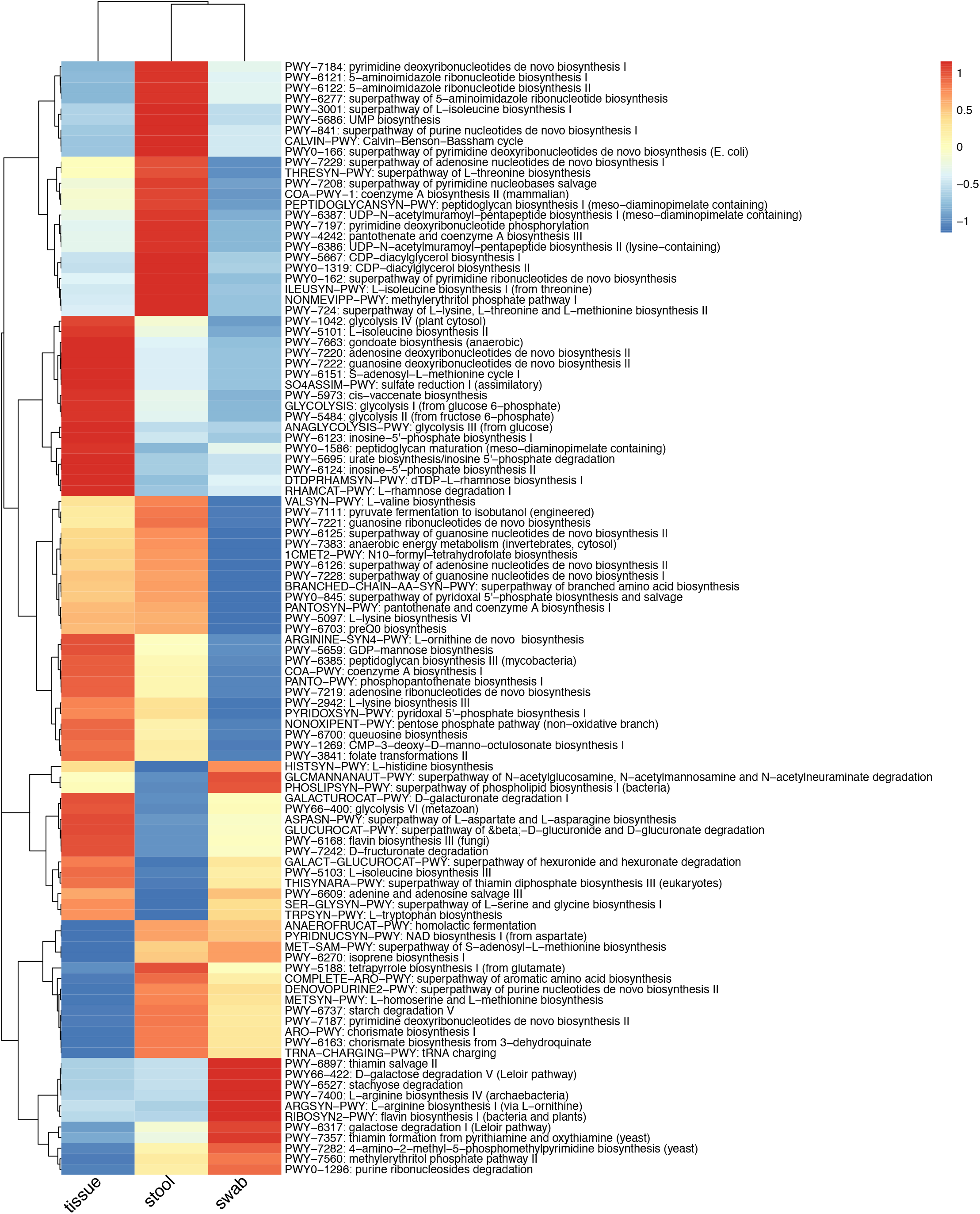
Heatmap of functional pathways that were significantly different between sample types (FDR<0.05). Keys indicate z-scores of averaged abundance.

### The impact of sample type on the associations between the taxonomic and functional profiles and host factors

We built separate models in each sample type to estimate whether the associations with taxonomic composition were consistent across sample types for the host factors age, sex, BMI, antibiotics use and NSAIDs use. The associations between genera and host factors were very highly correlated between stool and swab samples (Fig. 5: left panels) with Spearman’s correlation coefficients of p-value vs. p-value ranging from 0.501 for BMI to 0.75 for sex. The associations between stool and mucosal tissue samples (Fig. 5: middle panels) were significantly correlated except for sex with a P-value cutoff of 0.05, while the associations between swab and mucosal tissue samples (Fig. 5: right panels) were significantly correlated for BMI, antibiotics use and NSAIDs use but not for age or sex.

**Fig. 5.**
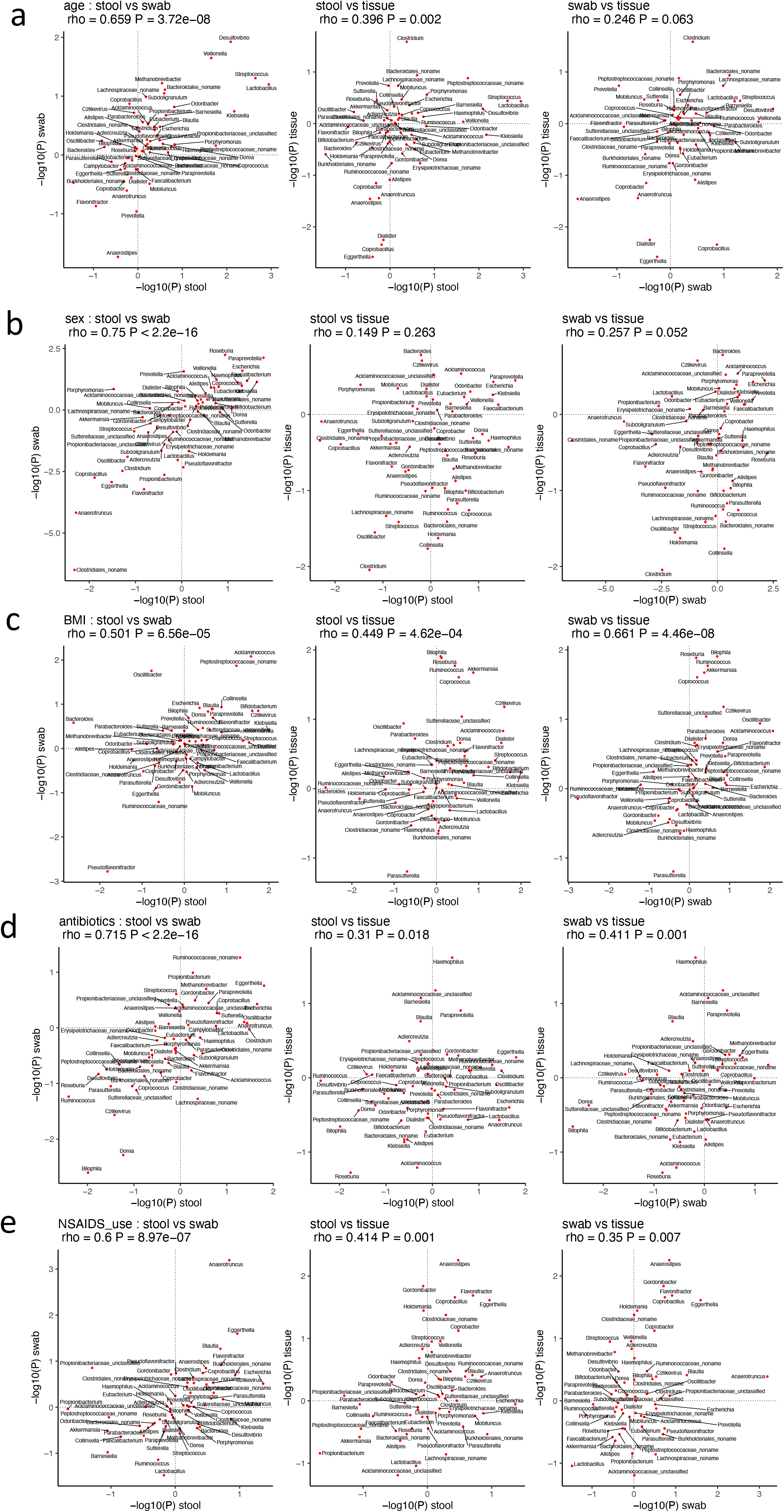
Correlations between the genus composition inference for age (a), sex (b), BMI (c), antibiotics use (d) and NSAIDs use (e) between pairwise sample types. The axes were the −log10 transformation of p-values from the model 2 described in methods

The same models were used for analyzing the robustness of the associations between pathways and host factors (Fig. 6). As was the case for taxa, the associations between pathways and host factors observed in stool and swab sample types were all highly positively correlated (Fig 6: left panels). However, comparisons between mucosal tissue and stool (Fig 6: middle panels) and swab (Fig 6: right panels) samples showed that the correlations were less consistent, including positive correlation with a smaller coefficient, negative correlation and no correlation. These observations were generally consistent when using ALDEx2 for statistical modeling instead of the linear models for both taxonomic composition and functional pathways that inference with stool and swab are more consistent than with mucosal tissue (Table S4 and S5).

**Fig 6.**
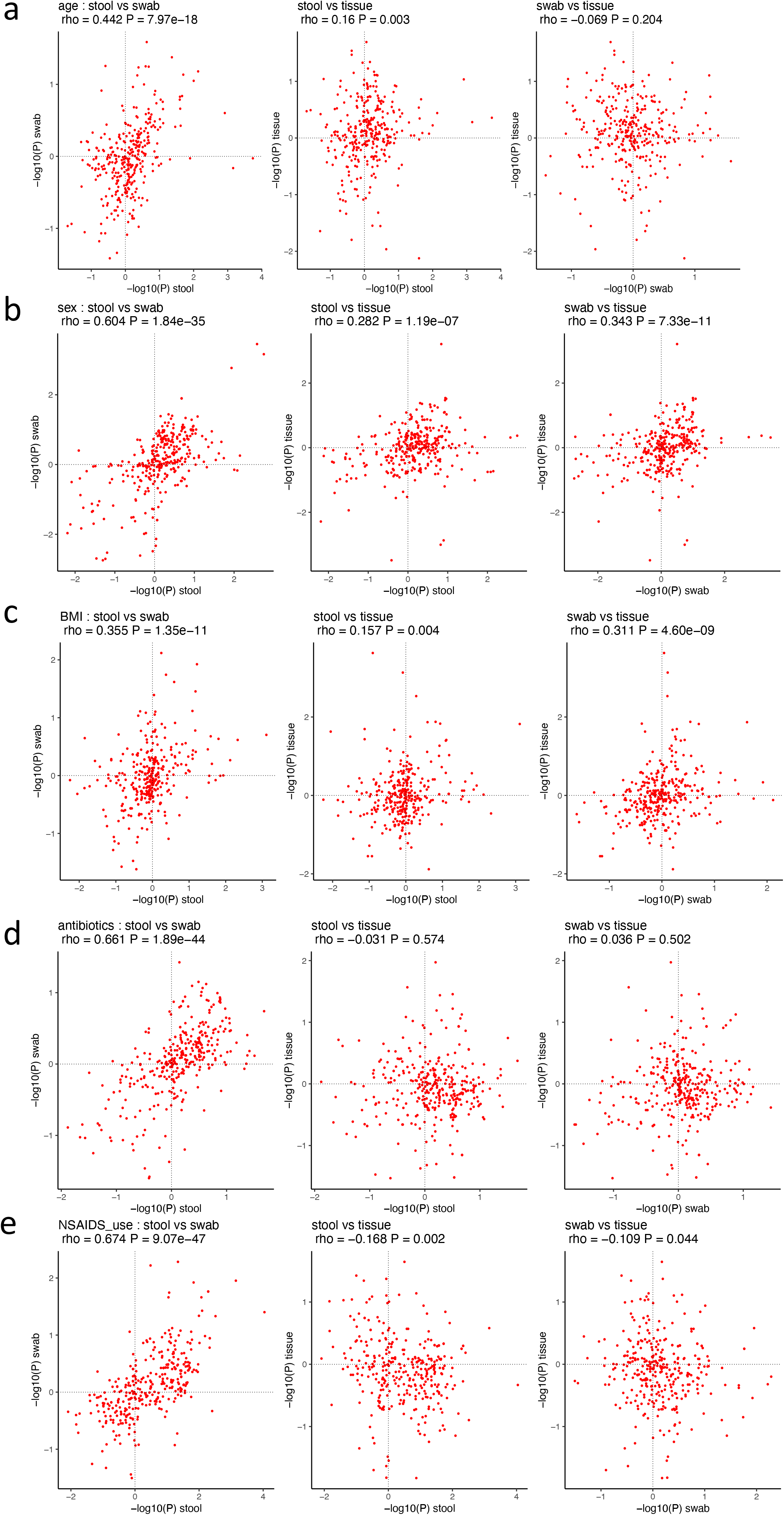
Correlations between the functional pathways inference for age (a), sex (b), BMI (c), antibiotics use (d), NSAIDs use (e) and between pairwise sample types. The axes were the –log10 transformation of p-values from the model 2 described in methods.

## Discussion

A better understanding of the associations between the human gut microbiome and disease is essential for developing potential early detection and intervention methods utilizing the microbiome. The stool, rectal swab and mucosal tissue biospecimen types we examined in this study sample microhabitats in which different microbial communities reside. With 1397 matched stool, rectal swab and mucosal tissue metagenomes for 240 participants, our dataset provided a great opportunity for analyzing the variations of these three matched biospecimens from the same participants. Unsurprisingly, we found that microbial taxonomic composition and functional pathways were different across the three biospecimen types, with the mucosal tissue metagenome more distinct from stool and swab. In general, the inference of host factor and microbiome associations were highly consistent between stool and rectal swab but not for mucosal tissue.

The mucosal tissue microbiome had lower alpha diversity and low abundance of most microbes, but was enriched in *Bacteroides, Subdoligranulum, Escherichia* and *Propionibacteriaceae*. *Bacteroides thetaiotaomicron, B. caccae, B. fragilis* and *B. vulgatus* are well known mucin degraders and rely on mucin and other host-derived glycans for colonization [15]. *Propionibacterium* (phylum Actinobacteria) and *Escherichia* (phylum Proteobacteria) were higher in mucosal tissue and swab compared to stool samples, which could be explained by their higher oxygen tolerance. The enrichment of Actinobacteria and Proteobacteria in the mucosa-associated microbiota has been reported in correlation with the intestinal radial colonic oxygen gradient that influences microbiota composition based on their ability to tolerate the oxidative stress [16]. The higher alpha diversity in the rectal swab microbiome compared to the stool and mucosal tissue microbiome is consistent with our previous study [8] and could be explained by swab sampling from both luminal and mucosal microbes [9].

Similar to taxonomic composition, the functional pathways in stool and rectal swab samples were more similar to each other than mucosal tissue samples. The number of sequencing reads from the mucosal tissue was smaller compared to stool and rectal swab samples due to lower microbial biomass and a higher percentage of human genome DNA contamination (Fig. S1). This could contribute to the observed lower taxonomic and functional diversity in mucosal tissue microbiome compared to stool and rectal swab samples. Compositional artifacts associated with reduced sequencing depth may therefore explain some of the differences we observed between mucosal tissue and stool and swab samples. These differences did persist even when using the compositionally aware pipeline ALDEx2, but no statistical approach can perfectly compensate for large differences in sequencing depth. ALDEx2 does not allow for inclusion of covariates or adjusting for random effects from the same subject and that might explain the differences between the ALDEx2 and linear models. Future research will be needed to explore how much of the differences between mucosal tissues and stool and swab in both community and gene composition and inference can be explained by these compositional differences.

Stool, swab and mucosal microbiota were enriched for different pathways, reflecting the niche adaption of different microbial communities. Mucosal microbiota was relatively enriched for pathways related to glycolysis and biosynthesis pathways involved in the generation of amino acid L−isoleucine, nucleosides adenosine, guanosine and inosine, and fatty acids gondoate and *cis*-vaccenate (one of the major unsaturated fatty acids, responsible for membrane phospholipid homeostasis in bacteria[17]). The stool and rectal swab microbiomes differed in the pathway related to peptidoglycan, CDP−diacylglycerol, UDP−N−acetylmuramoyl−pentapeptide, galactose, stachyose, L−arginine, purine and pyrimidine. Because a large number of functional genes remained unexplored, future expansion of database could provide a better knowledge of the functional differences between these sample types.

In order to determine whether the biospecimen type influence the inference of associations between the gut microbiome and host factors, we analyzed microbial associations with age, sex, BMI, antibiotics and NSAIDs use in each of the three sample types. We found that inferences performed with stool and rectal swab samples were highly correlated with each other for both taxonomic composition and functional pathways, while inference with mucosal tissue was more distinct especially for functional pathways. The relatively poor consistency between the mucosal tissue microbiome and the stool and rectal swab microbiome potentially reflects the niche differences that affect microbial interactions with the environment. It is also possible that the mucus barrier between the mucosal tissue and the lumen makes the mucosal tissue microbiome more sensitive to some host changes that were reflected in the mucosal tissues. For example, a previous study reported that the excessive secretion of mucus glycan could lead to the increase of *Akkermansia* and *Bacteroides* abundance in mucosal tissue but was only extended to stool with an altered mucus barrier [7]. As is the case for comparisons of relative abundance, models of inference are also sensitive to compositional artifacts associated with sequencing depth, although in our study comparisons based on ALDEx2 yielded broadly similar results to comparisons based on compositionally naïve mixed linear models.

We note that this study was conducted in individuals with a history of colorectal polyps, so the conclusions may not be generalizable to individuals without a history of polyps. However, all the participants were polyp-free when biospecimens were collected. Our work represents the largest study to date to explicitly compare these sample types and should provide a useful guide to investigators in the design and interpretation of human studies of the gut microbiota.

## Conclusion

Our study shows that the stool, swab and mucosal tissue microbiota are of different taxonomic and functional profiles, but the stool and swab microbiota are generally more similar compared to that of mucosal tissue. When analyzing the associations between microbiota and host factors of age, sex, BMI, antibiotics or NSAIDs use in each sample type, the inference on stool and swab samples were also more consistent than the inference on mucosal samples. Our study suggests that not only the taxonomic and functional profiles varied by sample types but the inference on their associations with host factors were depending on the sample type as well.

## Acknowledgements

The authors thank the research staff and investigators who have contributed to the Personalized Prevention of Colorectal Cancer Trial, the study participants who contributed their time and biospecimens for research, and Dr. Jay Fowke for sharing his protocol for rectal biopsy collection. This study was supported by grants R03CA183019 (MJS), R01CA149633 (QD and CY), and R01DK110166 (QD and MJS), as well as the Ingram Cancer Center Endowment Fund. Data collection, sample storage and processing for this study were partially conducted by the Survey and Biospecimen Shared Resource, which is supported in part by P30CA68485. Clinical visits to the Vanderbilt Clinical Research Center were supported in part by the Vanderbilt CTSA grant UL1 RR024975 from NCRR/NIH. The UNC Microbiome Core is supported in part by P30 DK034987 Center for Gastrointestinal Biology and Disease (CGIBD) and P30 DK056350 UNC Nutrition Obesity Research Center (NORC). The parent study data were stored in Research Electronic Data Capture (REDCap) and data analyses (VR12960) were supported in part by the Vanderbilt Institute for Clinical and Translational Research (UL1TR000445). The content of this paper is solely the responsibility of the authors and does not necessarily represent the official views of the National Cancer Institute or the National Institutes of Health.

## Author Contributions

QD, CY, MJS and AAF contributed to study conception, design, and supervision. XZ, HJM, RMN, DLS, MAAP, QD and MJS contributed to acquisition of data. XZ and MJS provided administrative, technical, or material support. SS, XZ, HX, AS, IB, CY, DQ, MJS and AAF contributed to analysis and interpretation of data. All authors contributed to writing, review, and/or revision of the manuscript and approved the final manuscript.

## Data sharing statement

The metagenomes sequences analyzed in this study are available at NBCI with accession ID PRJNA693850. Scripts used in this study are available at https://github.com/ssun6/StoolSwabTissue.

