## Supplementary material for "On the robustness of inference of association with the gut microbiota in stool, swab and mucosal tissue samples": SI.pdf

### Supplementary Table

Table S1. PERMANOVA test of the association between sample types and taxonomic composition of microbial metagenomes from phylum to species level.

| | Stool, swab and tissue | | Stool and swab | |
|---|---|---|---|---|
| | $R^2$ | P | $R^2$ | P |
| phylum | 0.265 | 0.001 | 0.035 | 0.001 |
| class | 0.404 | 0.001 | 0.041 | 0.001 |
| order | 0.397 | 0.001 | 0.056 | 0.001 |
| family | 0.351 | 0.001 | 0.053 | 0.001 |
| genus | 0.316 | 0.001 | 0.050 | 0.001 |
| species | 0.256 | 0.001 | 0.040 | 0.001 |

Table S4. Correlation of the associations between taxonomic composition and host factors in stool, swab and tissue samples estimated with ALDEx2.

| | **stool vs swab** | | **stool vs tissue** | | **swab vs tissue** | |
|---|---|---|---|---|---|---|
| | rho | P | rho | P | rho | P |
| age | 0.193 | 0.025 | -0.037 | 0.731 | -0.083 | 0.430 |
| BMI | -0.180 | 0.038 | 0.020 | 0.849 | 0.006 | 0.955 |
| sex | 0.427 | 3.55E-07 | 0.001 | 0.991 | 0.037 | 0.726 |
| NSAIDS_use | 0.206 | 0.017 | 0.022 | 0.840 | 0.045 | 0.668 |
| antibiotics | 0.326 | 1.70E-04 | -0.070 | 0.515 | -0.174 | 0.099 |

Table S5. Correlation of the associations between functional pathways and host factors in stool, swab and tissue samples estimated with ALDEx2.

| | **stool vs swab** | | **stool vs tissue** | | **swab vs tissue** | |
|---|---|---|---|---|---|---|
| | rho | P | rho | P | rho | P |
| age | 0.105 | 0.026 | -0.112 | 0.031 | -0.012 | 0.814 |
| BMI | -0.028 | 0.549 | 0.138 | 0.008 | -0.025 | 0.626 |
| sex | 0.376 | 2.22E-16 | 0.008 | 0.883 | -0.033 | 0.522 |
| NSAIDS_use | 0.41 | 2.18E-19 | 0.15 | 0.004 | -0.028 | 0.589 |
| antibiotics | 0.308 | 3.63E-11 | 0.062 | 0.232 | 0.117 | 0.024 |

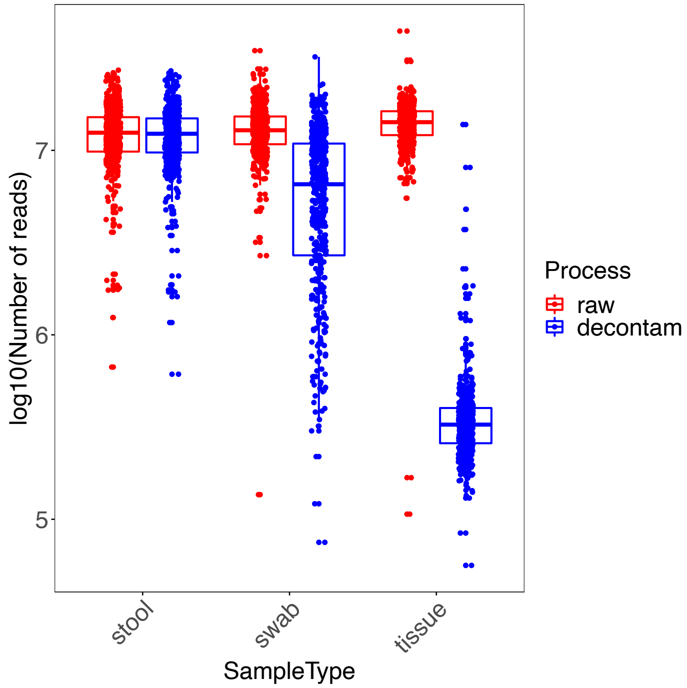


Fig. S1. Number of sequencing reads before and after removing human genome contamination in each sample type.

a

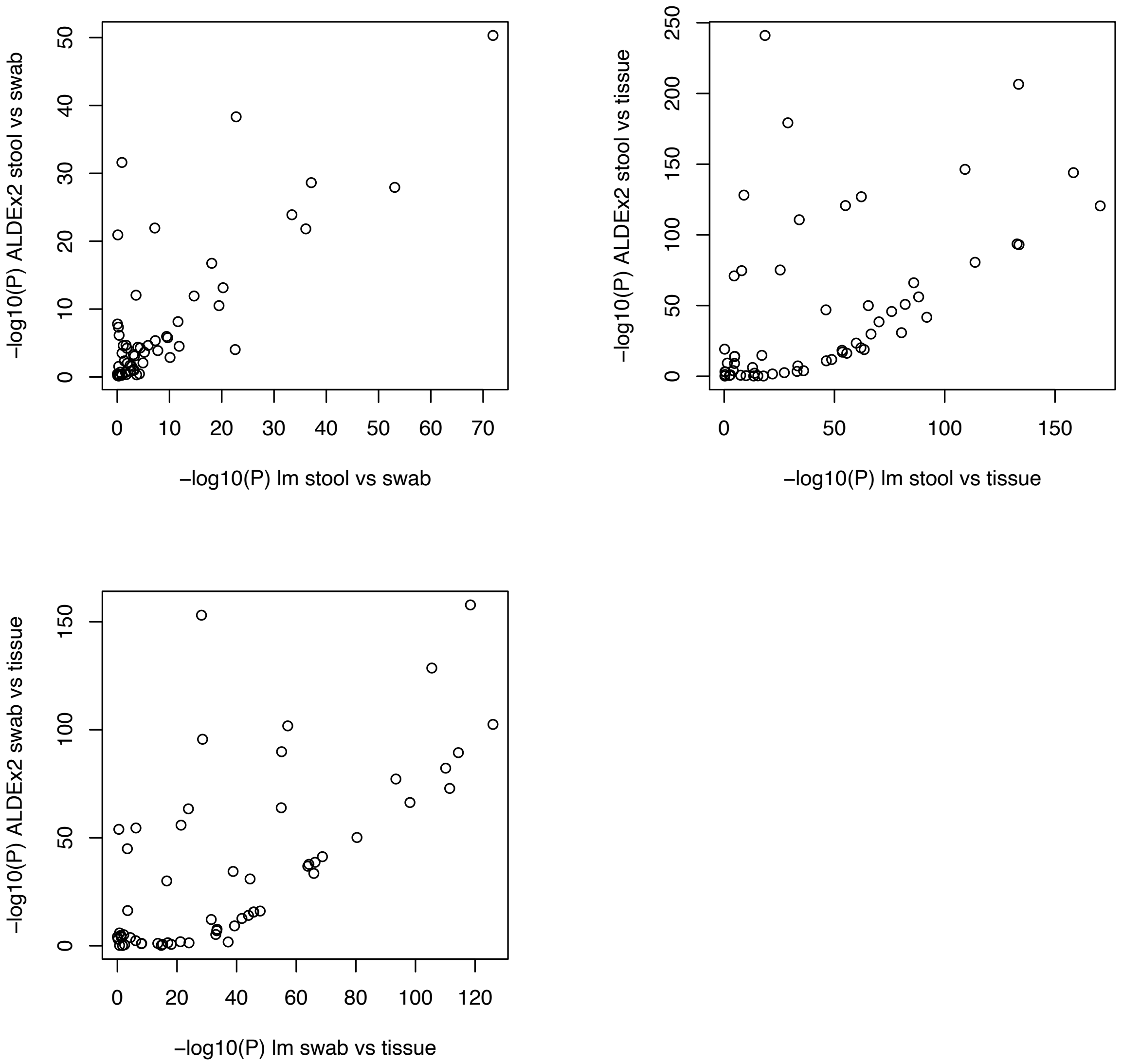

−log10(P) ALDEx2 stool vs swab
−log10(P) lm stool vs swab
−log10(P) ALDEx2 stool vs tissue
−log10(P) lm stool vs tissue
−log10(P) ALDEx2 swab vs tissue
−log10(P) lm swab vs tissue

b

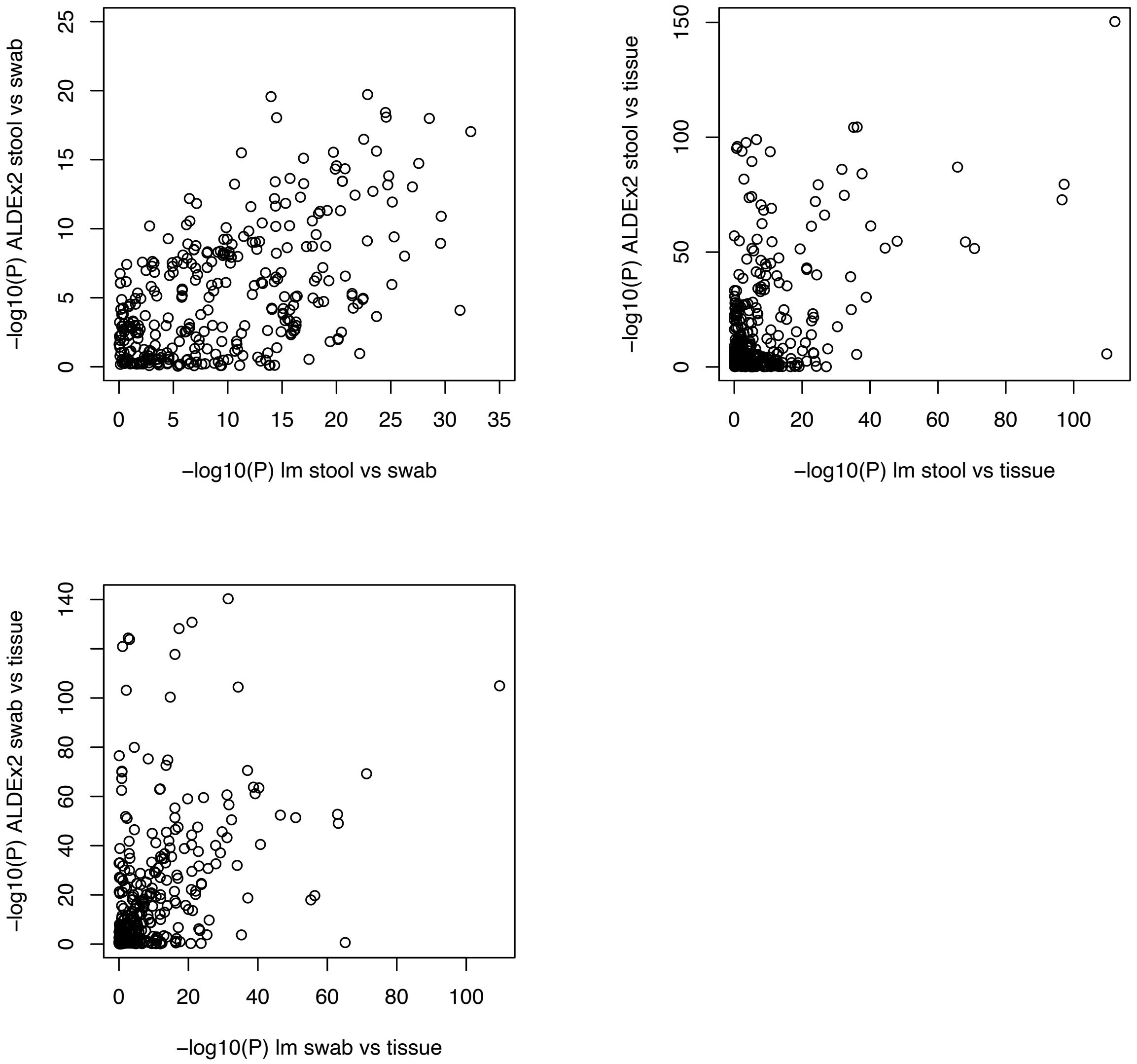


Fig. S2. Comparison of P values from ALDEx2 and mixed effects linear models for each pair of sample types for genus (a) and pathway abundance (b).
